# Roughness of a transmembrane peptide reduces lipid membrane dynamics

**DOI:** 10.1101/093377

**Authors:** Marie Olšinová, Piotr Jurkiewicz, Jan Sýkora, Ján Sabó, Martin Hof, Lukasz Cwiklik, Marek Cebecauer

## Abstract

Transmembrane domains integrate proteins into cellular membranes and support their function. The capacity of these prevalently α-helical structures in mammals to influence membrane properties is poorly understood. Combining experiments with molecular dynamics simulations, we provide evidence that helical transmembrane peptides with their rough surface reduce lateral mobility of membrane constituents. The molecular mechanism involves trapping of lipid acyl chains on the rough surface and segregation of cholesterol from the vicinity of peptides. The observations are supported by our toy model indicating strong effect of rough objects on membrane dynamics. Herein described effect has implications for the organization and function of biological membranes, especially the plasma membrane with high cholesterol content.

---

Membrane proteins influence many vital functions of living cells. They can constitute up to 50% of membrane total mass.^1^ Proteins associate with membranes peripherally via protein-protein or protein-lipid interactions and use lipid anchors for more stable attachment, but can also be fully integrated by spanning a membrane. Such integration requires the presence of one or multiple transmembrane domains (TMDs) in protein structure to overcome the hydrophobicity of lipid membranes. Direct contact of TMDs with lipids can influence membrane organization and dynamics but the molecular mechanisms responsible for this effect are still poorly understood.

TMDs of integral proteins can be considered as short peptides composed of 15-30, prevalently non-polar amino acids. In mammals, they form mainly α-helix stably integrated into the hydrophobic core of lipid membranes. In addition to membrane integration and stabilization, TMDs can affect ternary/quaternary structure and oligomerization of proteins.^2,3^ TMDs are often represented by smooth cylinders in literature (Fig. 1a). Sometimes such smooth cylinders are even associated with lipids.^4^ However, stable helical peptides formed by native amino acids are intrinsically rough (Fig. 1b and Supplementary Fig. 1). Indeed, a rough surface decorates all TMDs of proteins characterized so far by crystallography or NMR at sufficient resolution (< 4.5 Å; PDBTM database - <u>pdbtm.enzym.hu</u>).^5^

**Figure 1.**
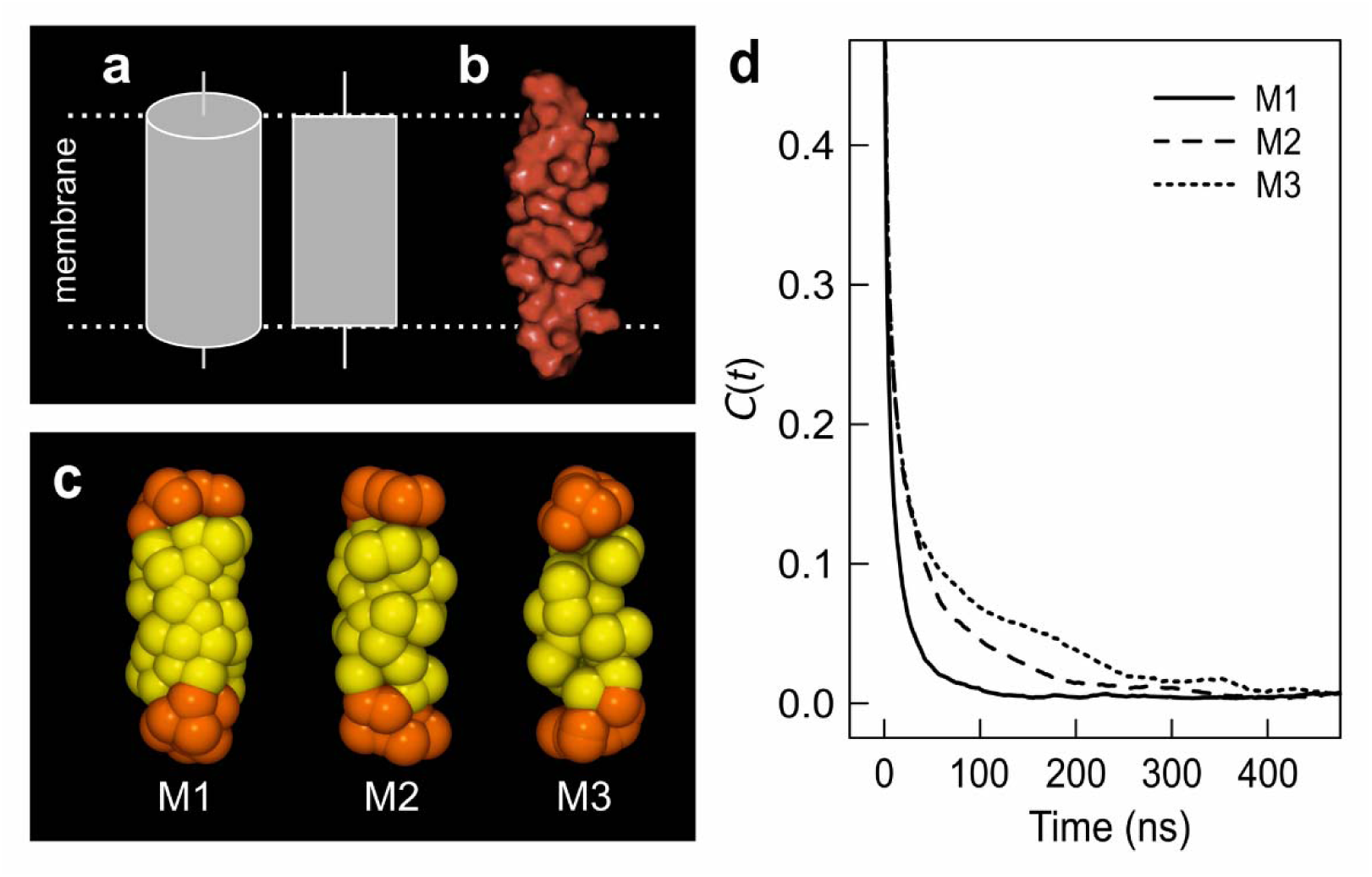
Rough surface interferes with membrane dynamics. **a.** Common styles of schematic illustration of TMDs as smooth cylinders, **b.** On the contrary, helical peptides have inherently rough surface formed by amino acid side chains. **c.** Toy models (M1-M3) of cylindric-like structures generated using coarse grain force field. **d.** The autocorrelation curves for contacts between lipid tails and the surface of model structures M1-M3 embedded in DOPC membrane with increasing roughness.

Diffusion of polymers^6^ and peptides^7^ is known to be slowed down near surfaces with nanoscopic roughness. Therefore, we were interested how cylinder-like objects with smooth versus rough surface influence the mobility of lipids and their acyl chains in membranes. The coarse grain toy models with varying surface roughness were generated (Fig. 1c) and embedded in dioleoyl phosphatidylcholine (DOPC) membranes. The autocorrelation data acquired from molecular dynamics (MD) simulations indicate slower lipid-surface contact dynamics for rough surfaces (Models M2 and M3) compared to the smooth ones (Fig. 1d). In this model, acyl chains are entrapped in the grooves of the rough objects which hinders their mobility (Fig. 1c and Supplementary Fig. 2 (new)). Virtually no trapping of acyl chains was detected at the smooth surface (Fig. 1d and Supplementary Fig. 2; Model M1). This model indicates that objects with rough surface embedded in membranes reduce mobility of its components.

To experimentally verify this prediction, we used simple proteo-lipid model membranes in the form of large and giant unilamellar vesicles, LUVs and GUVs, respectively (Supplementary Fig 3 and Supplementary Discussion, Section 1). Their composition was adjusted to selectively investigate a direct impact of transmembrane peptide on a membrane in the absence of previously described effects which are independent of its roughness; e.g. the impact of obstacles (crowding), hydrophobic mismatch and formation of clusters or lipid domains.^8^-^11^ The lipid with low melting temperature (DOPC, T_m_ = -18.3°C)^12^ was selected to preserve the fluid character of membranes under all tested conditions and avoid formation of lipid domains.^13^ The synthetic, highly purified, transmembrane peptide (LW21: GLLDSKK<u>WWLLLLLLLLALLLLLLLLWW</u>KKFSRS) represents a stable, monomeric structure with a rough surface in lipid membranes (Supplementary Fig. 4).^14^ Since proteins account approximately for 2-3 mol% of cellular membranes (see Supplementary Discussion, Section 2), we examined model membranes containing 0-3 mol% of LW21 peptide. Calibration-free z-scan FCS technique^15^ was used to measure lateral diffusivity of fluorescent lipid tracer, DiD, and fluorescently-labelled LW21 embedded in membranes. In Fig. 2a, we demonstrate that increasing the concentration of monomeric a-helical peptide reduced lateral diffusivity of both lipid tracer and labelled peptide (grey and red lines, respectively). At the highest tested peptide concentration, 3 mol%, diffusivity of both molecules was reduced by approximately 35% in DOPC membranes. In agreement with the literature,^9^ lipid molecules diffused somewhat faster than peptides at all tested peptide concentrations. To better mimic cell membranes’,^16^ the experiments were also performed in presence of 25 mol% of cholesterol. The impact of the peptide on the lateral diffusion of membrane components was more pronounced in the presence of cholesterol than in its absence. At the highest peptide concentration, 3 mol%, we observed 2-3 fold decrease in the diffusion coefficients for both tested molecules (Fig 2a). Quantitative analysis (D_25_/D_0_ values) supports the enhanced effect of peptide on the mobility of membrane components in the presence of cholesterol compared to its absence (Supplementary Table 1). Lateral diffusion of lipids and peptides was indistinguishable in membranes with cholesterol (Fig 2a; blue and yellow lines).

**Figure 2.**
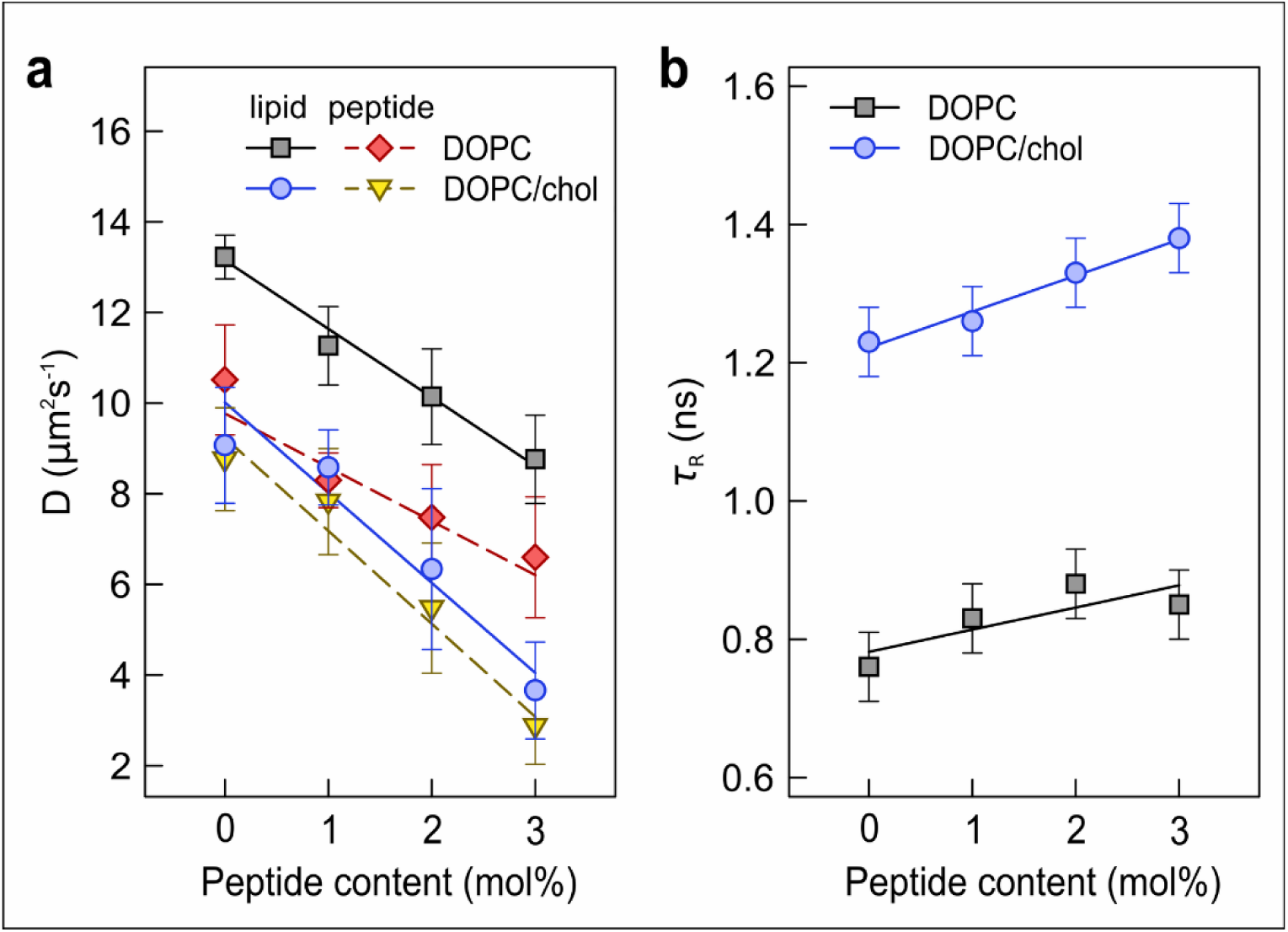
Impeded local viscosity and lateral diffusion in membranes with peptide. **a.** Lateral diffusion coefficients of lipid tracer (DiD; black and blue lines) and fluorescently labelled LW21 peptide (red and yellow lines) were measured in GUVs composed of DOPC (full lines) and DOPC:cholesterol (3:1; dashed lines), in the presence of increasing concentration of unlabeled peptide. Each presented diffusion coefficient (D) was measured for at least 10 vesicles in three independent experiments using callibration-free z-scan FCS (fluorescence correlation spectroscopy) technique. Error bars indicate standard deviations (SD). **b.** Local lipid mobility (viscosity) as a function of increasing peptide concentration was determined in the absence (black squares) or presence (blue circles) of 25 mol% cholesterol using Laurdan fluorescent probe by TRES. Error bars represent intrinsic uncertainty of the method.

Membrane dynamics was further evaluated at the nanoscale using environment-sensitive fluorescent probes. First, time-resolved emission spectra (TRES) of Laurdan were measured. The speed of Laurdan TRES red-shift, represented by the relaxation time τ_R_, reports on the local mobility of lipid carbonyls located at the interface of the hydrophobic and hydrophilic parts of the lipid membrane (Supplementary Fig. 5). Significant increase of τ_r_ as a function of peptide content in DOPC membrane entails that LW21 hinders mobility of lipid carbonyls (Fig. 2b). This hindrance was slightly stronger in the presence of cholesterol compared to its absence (Fig. 2b; linear regression slope of 0.05 versus 0.03, respectively). Second, time-resolved anisotropy of diphenylhexatriene was measured to characterize the order and mobility of lipid tails creating the hydrophobic interior of the membrane. The obtained results (Table 1 and Supplementary Discussion, Section 3) show restriction in DPH rotation manifested in the elevated DPH order parameter in the presence of peptide. This effect was preserved in cholesterol-containing membranes, which agrees with the previously published data.^17^

**Table 1.** Order parameter S of DPH in membranes with the LW21 peptide.

| Membrane (LUV) | S (DOPC) | S (DOPC/cholesterol) |
| --- | --- | --- |
| DOPC | $0.19 \pm 0.06$ | $0.43 \pm 0.01$ |
| DOPC + 3% LW21 | $0.28 \pm 0.04$ | $0.51 \pm 0.05$ |
| DOPC + 10%<br>LW21 | $0.39 \pm 0.02$ | $0.55 \pm 0.08$ |

To verify that the rough surface is responsible for the reduced membrane dynamics observed in our experiments, we have performed fully atomistic MD simulations of the peptide in the lipid bilayers composed of DOPC both without and with cholesterol (Supplementary Movies 1-3). We confirm considerably impeded mobility of lipids close to LW21 peptide (Supplementary Fig.6). In agreement with our model (Fig. 1), we observed that this is caused by the trapping of acyl chains of annular lipids in the grooves of the rough surface of the peptide, which is formed by amino acid side chains (Fig 3a,b). Such lipid-peptide contacts are non-specific and exhibit substantial stability (Supplementary Figs. 7 and 8). On the contrary, cholesterol was excluded from the annulus, thus exhibiting fewer contacts with the peptide compared to phospholipids (Fig. 3c-e and Supplementary Fig 9). This is probably caused by the incompatibility between the planar shape of cholesterol molecule and the roughness of transmembrane peptides (ref.^18^, Supplementary Fig. 10 and Supplementary Discussion, Section 4). Our findings demonstrate that the mobility of lipids is affected not only by large and multispanning proteins^19,20^, as reported in recent computational studies, but also by a single transmembrane domain due to its rough surface.

**Figure 3.**
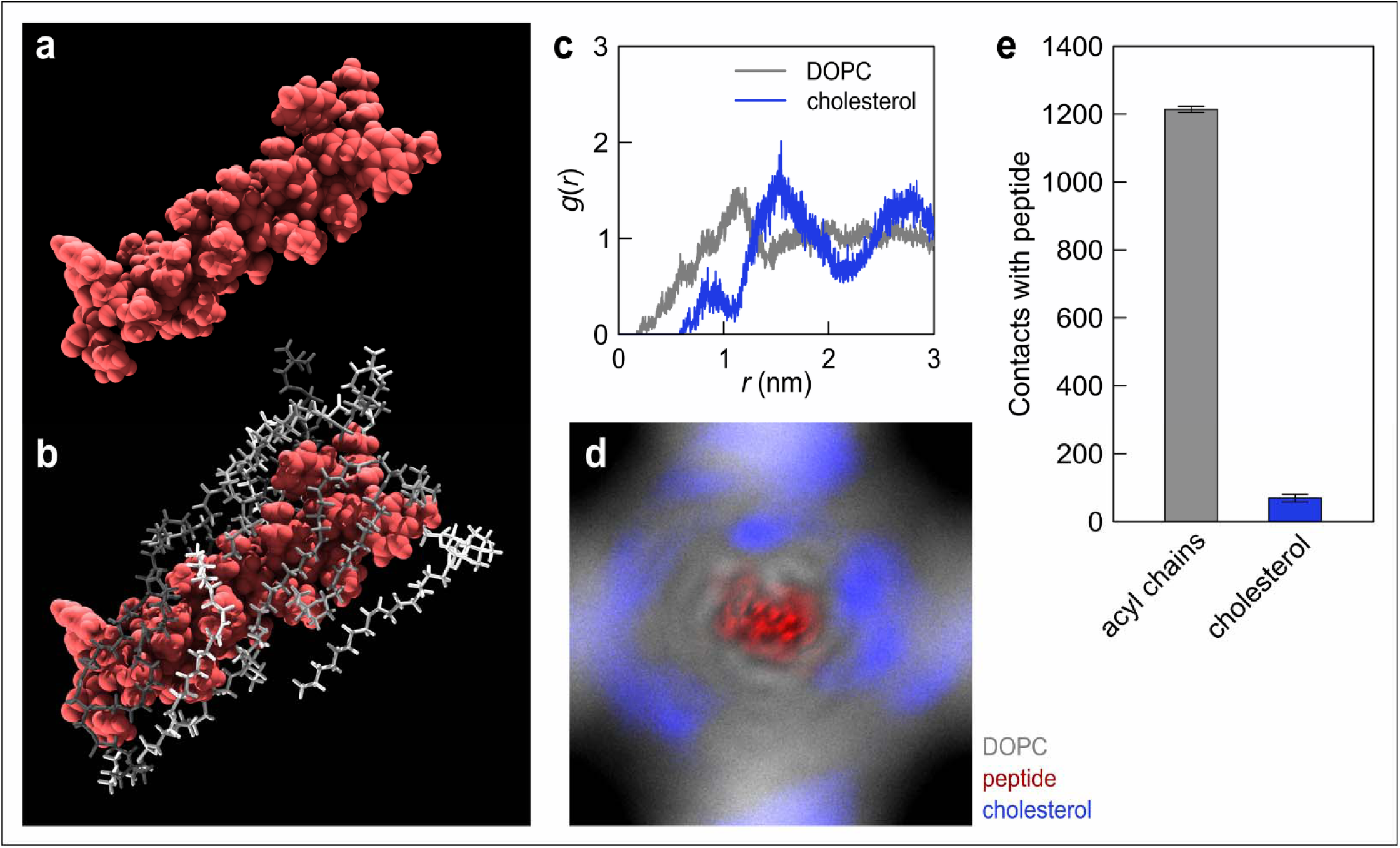
Non-specific lipid acyl chain trapping on the rough surface of LW21 peptide and cholesterol segregation – MD simulations. **a** and **b.** A typical snapshot from MD simulation of the peptide in DOPC bilayer indicating trapping of lipid acyl chains in the grooves formed by peptide side chains. Peptide surface (a) is shown in red. Interacting lipids are shown in different shades of grey using licorice representation (b). Non-interacting lipids and water were removed for clarity. **c.** Pair correlation function [g(r)] of phospholipids and cholesterol from the centre of mass of the peptide. This function quantifies the probability of intermolecular distances between the peptide and lipids with respect to those in an ideally mixed system. **d.** Distribution map of cholesterol (blue) and phospholipids (grey) in membrane with peptide (red). The peptide was centered and rotations were removed by data postprocessing. **e.** Quantification of phospholipid and cholesterol contacts with the peptide calculated in MD simulations. Error bars represent error of the mean estimated by the block averaging method.

Mobility of membrane molecules is described in Saffmann-Delbruck hydrodynamic model (SD model)^21^ highlighting the effect of viscosity and, partially, the size of molecules. The variants of SD model for large immobile objects ^22^ or multispanners in heterogenous lipid membranes ^19^ considered proteins (or their TMDs) as smooth cylinders with the effective radius of > 4nm (Fig.1a). The calculated effect of such objects on lateral diffusion was negligible at low concentrations < 5 mol%. In nature, most of proteins possess a single TMD.^23^ Their effective intramembranous radius is comparable to the size of lipids (< 1 nm). Therefore, we have investigated the impact of integral proteins at low, physiologically-relevant concentrations (< 5 mol%) and demonstrate that transmembrane helical peptide (a model of simplest TMD) reduces diffusivity of membrane components by exposing its rough surface to the surrounding lipids. This is in agreement with our toy model (Fig. 1) that objects with rough surface have stronger effect on membranes compared to the smooth ones. Rough surface causes lipid acyl chains trapping and, as a consequence, reduces lateral diffusion of membrane molecules. In ternary membranes, the effect is further escalated by segregation of planar molecule, cholesterol, from the rough surface. Importantly, this effect is observed in the absence of hydrophobic mismatch, detectable domains and the presence of large and immobile obstacles or peptide aggregates, thus supporting a direct effect of proteins with rough TMDs on lipid membranes. It is also not restricted to the peptide tested in this work since trapping of lipid acyl chains in the grooves of the surface of a membrane protein was recently observed in crystals of aquaporin-0^24^ and Na^+^K^+^ pump.^4^ A rough surface of transmembrane domains, with potential for acyl chain trapping, was shown for a number of proteins (for example PDB ID: 5EE7^25^ or 2NA8^26^, Supplementary Fig. 1 and Supplementary Discussion, Section 5).

The physical effect of the TMD roughness on membrane dynamics and organization demonstrated in this work may have previously unexpected consequences for cell membranes and associated processes (see also Supplementary Discussion, Section 6). For example, it implies that reorganization of proteins (e.g. receptor clustering by ligands) might cause a local reduction in the mobility of membrane molecules. Inverse correlation between protein density and lateral diffusion that has been observed in both model and cell membranes in past might not be exclusively due to crowding.^27,28^ In addition, a tendency of cholesterol to segregate from TMDs might help formation and/or stabilization of cholesterol-enriched, protein-low domains. Such changes will locally affect intermolecular interactions or reaction kinetics of cellular processes associated with membranes.

## Methods

Methods and any associated references are available in the online version of the paper.

## Acknowledgements

We would like to thank Anthony I. Magee for stimulating this project and Peter Kapusta for technical assistance. This work was funded by Czech Science Foundation (M.C.: P305-11-0459, 15-06989S; L.C.: 15-14292S). M.C. would like to acknowledge Purkyne Fellowship and M.H. Praemium Academiae Award, both from the Czech Academy of Sciences.

## Author contributions

M.O. and M.C. conceived the project; M.O., P.J., J.Sy., L.C. and M.H. designed the experiments, M.O., P.J., J.Sa. and L.C. performed the experiments and M.O., P.J., L.C. and M.C. written and edited the paper.

## Additional information

Any supplementary information is available in the online version of the paper. Correspondence should be addressed to M.C.

## Online Methods

### Chemicals and peptides

All chemicals and organic solvents were purchased from Sigma-Aldrich and Merck, or otherwise stated. Lipids 1,2-dimyristelaidoyl-sn-glycero-3-phosphocholine (14:1 PC), 1,2-dioleoyl-sn-glycero-3-phosphocholine (DOPC; 18:1 PC), 1,2-dierucoyl-sn-glycero-3-phosphocholine (22:1 PC), 1,2-dipalmitoyl-*sn*-glycero-3-phosphoethanolamine-N-(cap biotinyl) (biotinylated-DPPE), 1,2-dioleoyl-sn-glycero-3-phosphoethanolamine (DOPE) and cholesterol (from ovine wool) were purchased from Avanti Polar Lipids, Inc. (Alabaster, AL, USA), streptavidin from IBA (Goettingen, Germany), and 2,2,2-trifluoroethanol (TFE) from Alfa Aesar (Karlsruhe, Germany). Fluorescent probes Dil.C_18_ (DiD) and 1,6-diphenyl-1,3,5-hexatriene (DPH) were purchased from Sigma-Aldrich, 6-lauroyl-2-dimethylaminonaphthalene (Laurdan) from Molecular Probes (Eugene, OR, USA), and Atto 488-maleimide from Atto-Tec (Siegen, Germany). Atto633-DOPE was prepared in our laboratory by coupling of Atto633 NHS ester (Atto-Tec) to the amine of DOPE followed by size-exclusion chromatography.

Peptide LW21 ((MW 4119, GLLDS<u>KKWWLLLLLLLLALLLLLLLLWWKK</u>FSRS) and its fluorescently labelled variant Atto488-LW21 (MW 4933 Atto488-CGLLDSKKWWLLLLLLLLALLLLLLLLWWKKFSRS) were custom synthesized by VIDIA (Prague, Czech Republic). The identity and purity of the products (> 92%) were confirmed by mass spectrometry and analytical H P LC. The sequence of LW21 peptide contains 21 hydrophobic residues flanked by two lysine residues of the original LW peptide (underlined; ref. ^1,2^) and a native sequence of the N- and C-terminal membrane proximal motifs from human TCRζ (five N- and four C-terminal residues). Solutions of 100 μM LW21, 100 μM C_2_-LW21 and 1 μM Atto488-LW21 peptides were freshly prepared in TFE for each experiment. Dimeric peptide C_2_-LW21 (MW 8579, (CGLLDPKKWWLLLLLLLLALLLLLLLLWWKKFSRS)_2_; Biomatik, Wilmington, USA) was obtained by cysteine oxidation of monomeric peptide. The efficiency of dimerization (> 95%) was confirmed by mass spectrometry and HPLC. Synthetic melittin (MW 2845, GIGAVLKVLTTGLPALISWIKRKRQQ) was obtained from Sigma–Aldrich.

### Preparation of model membranes (vesicles)

For vesicle preparation, lipids (DOPC and cholesterol) were dissolved in chloroform, fluorescent probes (DPH and Laurdan) in methanol and unlabeled LW21, C_2_-LW21 and Atto488-LW21 peptides were dissolved in TFE. **LUV** (large unilamellar vesicle) suspension was prepared as described below. Lipids, peptide and fluorescent probes (probe:lipid molar ratio = 1:100) were mixed in the desired ratio in a glass tube. Organic solvents were evaporated under a stream of nitrogen while continuously incubated in a water bath (ambient temperature) and then kept under vacuum for at least 1 hour. The dry lipid film was hydrated in heated 105 mOsm/kg glucose buffer (40°C, ~75 mM glucose, 10 mM HEPES, 10 mM NaCI and phi = 7.4) and vortexed for 5 minutes. The cycles of heating and vortexing were repeated until the lipids and peptides were suspended in the solution. To facilitate the detachment of peptides and lipids from the surface during the vortexing, glass beads (extensively cleaned; 2 mm in diameter) were added to the tube. LUVs were obtained by extrusion of lipid and peptide suspension through a polycarbonate membrane with an effective pore diameter of 100 nm (Avestin, Ottawa, Canada) by 50 passages at ambient temperature. Monomeric control for anisotropy was prepared by adding aqueous solution of melittin peptide which spontaneously integrates into lipid membranes, to the suspension of DOPC LUVs.

**GUVs** (giant unilamellar vesicles) were prepared by mixing lipids, peptides and probes in a desired ratio in a glass vial (2 ml) - altogether 100 nmol of all lipid species. For fluorescence correlation spectroscopy (FCS) experiments, GUVs contained DOPC, 0 or 25 mol% of cholesterol, 0-3 mol% of LW21 peptide and 2 mol% biotinylated-DPPE for the immobilization of vesicles at the BSA-biotin/streptavidin-coated glass coverslips. In addition, fluorescently labelled peptide Atto488-LW21 (peptide:lipid = 1:25 000) and lipid tracer, DiD (probe:lipid = 1:100 000), were added to lipid-peptide mixtures. For peptide incorporation studies, lipids of different acyl chain length were used (14:1 PC, 18:1 PC (DOPC), and 22:1 PC) and fluorescently labelled species were present in a higher content (Atto488-LW 1:1000 and fluorescent lipid, Atto633-DOPE, 1:20000). Otherwise the composition was identical to the GUVs used for FCS. GUVs were prepared by electroformation according to Stockl and co-workers.^3^ Briefly, lipids, peptides and fluorescent probes in organic solvents were spread on two preheated titanium slides. Slides were kept for 1 hour under vacuum to evaporate any residual solvent. The slides were then stuck together by melting Parafilm. The formed chamber was filled with 105 mOsm/kg sucrose in water and heated to 40°C. The process of vesicle formation was induced by applying a sinusoidal altering voltage (10 Hz) starting at 150 mV (peak-to-peak amplitude) and gradually increasing every 2.5 min with a step of 50 mV up to 1.1 V; this voltage was kept for another 90 minutes. At the end, a frequency of 4 Hz and voltage of 1.3 V were applied for 30 minutes for the detachment of vesicles. Prepared GUVs were transferred into a LabTek 8-well chamber slide (ThermoScientific, Waltham, USA) coated with BSA-biotin/streptavidin. Landing of GUVs on the optical surface was enabled by changing the buffer in the chamber for 105 mOsm/kg glucose (which has the same osmolality as sucrose buffer but higher density). Within one hour, GUVs were firmly attached to the coated glass surface and ready for imaging.

### Lateral mobility measurements - z-scan FCS

FCS measurements of GUVs were performed on an inverted confocal fluorescence microscope, Olympus IX71 (Olympus, Hamburg, Germany), equipped with single-photon counting unit MicroTime 200 (PicoQuant, Berlin, Germany). Excitation lasers of 470 and 635 nm (LDH-P-C-470 and LDH-D-C-635; PicoQuant), each with 10 MHz frequency, were used to illuminate a sample through the water-immersion objective (1.2 NA, 60x) (Olympus). The laser intensity measured at the sample position reached 5 |lW and 0.5 p,W for 470 nm and 635 nm lasers, respectively. The lasers were pulsing in alternating mode in order to avoid artefacts caused by signal bleed-through. Fluorescence signal was gathered through the main dichroic (Z473/635, Chroma, Rockingham, VT), 50 p,m pinhole and guided to the emission dichroic mirror (620DCXR, Chroma) which splits the signal between the two single photon avalanche diodes using 515/50 and 685/50 band pass filters (Chroma). The measurements were performed at 25°C.

The information about membrane components’ mobility was obtained by employing fluorescence correlation spectroscopy (FCS, for the details see ref. ^4^). Autocorrelation function of the temporal fluorescence intensity fluctuations contains the information about the average number of particles *N* in the detection volume and characteristic time, *τ_D_*, the molecule spends on average inside the detection volume (diffusion time). *N* and *τ_D_* are obtained by fitting the autocorrelation function with *a priori* known model: eq. 1 for the lipid lateral mobility, eq. 2 for peptide lateral mobility. Eq. 2 takes into account the free dye not bound to the peptide (i.e. correction for the sample impurity originating from peptide labelling).

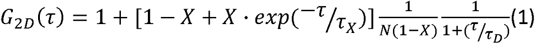

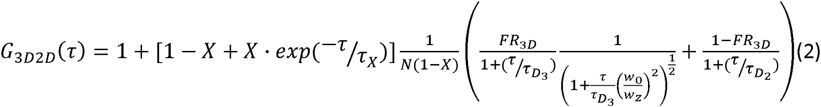

The *G_2D_*(*τ*) and *G_3D2D_(τ)* are calculated autocorrelation functions for the delay time, *τ, X* is an average fraction of molecules in dark state (due to intersystem crossing or cis/trans isomerization; typically set to zero for *G_3D2D_(τ)), τ_X_* represents dark state relaxation time, *FR_3D_* is a fraction of molecules moving in 3D space, *τ_D3_* and *τ_D2_* are characteristic diffusion times for molecules moving in 3D and 2D space, *w_0_* and *w_z_* are the lateral and radial dimensions of the confocal volume, respectively, obtained as the distance from and along the optical axis at which the intensity drops to *e^−2^*.

To overcome the difficulties of precise volume determination for the conversion of *τ_D_* determined by FCS to diffusion coefficient *D*, z-scan FCS variant was employed. Z-scan FCS represents a calibration-free method providing the diffusion coefficient and the radius of confocal volume in lateral plane.^5^ The z-scan FCS method is based on recording a set of measurements of the labelled membrane positioned differently in respect to the z-axis of the microscope. Intensity fluctuations are recorded in several consecutive z-positions separated by 200 nm with the top of GUV membrane placed ideally in the center of the z-stack. Analysis of calculated autocorrelation curves yields the set of *τ_D_.* Fitting the parabolic dependence of *τ_D_* the position along the microscope z-axis provides directly the lateral diffusion coefficient *D* (eq. 3).

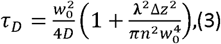

where *λ* is the wavelength of the excitation light, *Δz* is the distance between the sample position *z* and the position of the beam diameter minimum *z_0_, n* is refractive index of the medium.

### Determination of the peptide integration

Peptide integration studies were performed on an inverted confocal fluorescence microscope Olympus X71 with the same optical setup as for FCS experiments. The intensity image of each GUV was acquired by scanning the XY plane cross-section in the middle of the vesicle with a pixel size of 100 nm. Several line profiles of fluorescence intensity were calculated for each GUV; two maximal values along every line profile were determined. The average of these values was used to compare the peptide integration in membranes composed of different lipids (14:1 PC, 18:1 PC (DOPC), and 22:1 PC). At least 12 vesicles for each membrane composition were analyzed.

### Time-resolved emission spectra of the Laurdan probe

Time resolved emission spectra (TRES) were obtained by spectral reconstruction from the recorded steady state emission spectra and time resolved fluorescence decays. Steady-state fluorescence emission spectra (λ_EX_ = 376 nm) were recorded for the suspension of Laurdan-labelled LUVs with different peptide:lipid composition (lipid concentration: 1 mM) using Fluorolog-3 spectrofluorimeter (model FL3-11, JobinVvon Inc., Edison, NJ, USA) maintained at 25°C using a water-circulating thermostat. Fluorescence decays were collected by IBH 5000 U SPC equipment (Horiba Jobin Yvon) with a picosecond diode laser (IBH NanoLED-375LH, peak wavelength 376 nm, 1 MHz repetition rate). Decays were recorded at series of wavelengths spanning the steady-state emission spectrum (400–540 nm) in 10 nm steps. To eliminate scattered light, a 399 nm cut-off filter was applied. The signal was kept below 2% of the light source repetition rate, and the data were collected in 8192 channels (0.014ns per channel) until the peak value reached 5000 counts. Fluorescence decays and steady state spectra were used to reconstruct TRES, which were fitted with log-normal function to obtain the time evolution of their spectral maximum *v(t)*.^6^ Two main parameters were derived from *v(t):*

1) Overall emission shift *Δv*, which is directly proportional to the polarity of the dye microenvironment

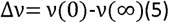

where *v(0)=* 23800 cm^−1^ (estimated using method of Fee and Maroncelli^7^) and *v(∞)* stands for the position of the TRES emitted from the fully relaxed state.

2) Relaxation time *τ_r_* refers to the viscosity of the dye microenvironment. τ_r_ reflects mobility of lipid moieties at the level of the carbonyl groups and peptide amino acid residues in the vicinity of Laurdan probe.

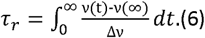

### Time-resolved fluorescence anisotropy

Fluorescence polarization experiments were performed using the same time resolved fluorescence spectrometer as for the TRES data. Measurements consisted of the recording of fluorescence decays with the 2 different orientations of the emission and excitation polarizers. The anisotropy decay r(t) was then calculated according to eq. (7):

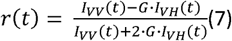

where *l_vv_* is the fluorescence decay measured with both excitation and emission polarized vertically, and *l_vh_* with the vertically polarized excitation and horizontally polarized emission. G-factor (*G*) was determined by acquiring fluorescence decays for a standard solution of free dye (10 μdM POPOP in ethanol or 40 μM tryptophan dissolved in PBS buffer) applying a tail matching method using FluoFit v.4.5 (PicoQuant). Anisotropy decay *r(t)* was fitted with single exponential

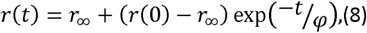

where *r*_∞_ is residual anisotropy, *r(0)* limiting anisotropy, and *ϕ* rotational correlation time.

Specifically, the anisotropy decays were measured either for DPH probe embedded in the membrane or for the tryptophans of LW21 peptides incorporated in the membrane. For the former set of experiments, the DPH-labelled LUV dispersion (lipid concentration: 0.5 mM) was excited by the picosecond diode laser 376 nm (IBH NanoLED-375LH, peak wavelength 376 nm, 1 MHz repetition rate). The emission monochromator was set to 466 nm. A 399 nm cut-off filter was applied to eliminate scattered light. The difference between the maxima of *l_vv_*, and *l_vh_* decays was set to 20000 counts. The anisotropy data were analyzed according to the “wobble in cone” model introduced by Kawato et al. (1977)^8^ and Kinosita et al. (1982)^9^. The S-order parameter (*S*) was calculated according to eq. 9:

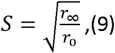

In the case of tryptophan anisotropy, dispersion of LUVs (0.5 mM lipid concentration) with indicated peptides was excited by 295 nm diode (Picoquant; PLS-8-2-257, 5 MHz repetition rate). The excitation light was guided through the polarizer directly into the sample and the emission monochromator was set to 350 nm. The difference between the maxima of *l_vv_* and *l_vh_* decays was set to 50000 counts. The data were collected in 4096 channels (0.028 ns per channel). The rate of Trp anisotropy decay determines the efficiency of the energy homo-transfer. A higher transfer efficiency is caused by the close proximity of the two helices (their tryptophan side chains), suggesting that a peptide forms dimers (or higher oligomers).

### Expression of LW21-GFP proteins in human cells

DNA plasmid for the expression of LW21-GFP was prepared in analogy with LW19-GFP used in our previous work.^10x^ Synthetic oligonucleotides FWD 5’-GAGCAGAAGCTTATCAGCGAAGAG and REV ATTAGATCTGCTGAACTTCTTCCACCACAAGAGAAGCAGAAGCAAGA-3’ were used as primers, and pXJ41-LW19-GFP as DNA template for polymerase chain reaction (PCR) amplification of LW21-GFP sequence. Full construct encoded 5’UTR and signal peptide from CD148, myc-tag and LW21 TMD linked to the EGFP via a short linker (see ref. ^10^ for more details). Our previous data indicate that signal sequence and c-myc tag do not influence protein sorting in tested cells.^10^

Jurkat cells were transiently transfected using Neon^®^ transfection system (Life Technologies). According the manufacturer’s instructions, 1μg of vector DNA per shot per 50 000 cells was used. Cells were imaged on poly-L-lysine-coated (to immobilize cells) glass-bottom 8-well chamber slides (Lab-Tek^®^, Thermo Scientific) supplemented with pre-heated, color-free RPMI-1640 medium. Images were taken using a laser scanning confocal microscope Leica SP5 TCS AOBS Tandem equipped with Leica HyD hybrid detector, 63x 1.3 NA glycerin immersion objective (Leica PLAN APO) and live cell support chamber. LAS AF image software (Leica Microsystems) was used for acquisition. Minor contrast/level adjustments were applied and images were processed for publishing by Fiji/ImageJ.^11^

### All-atom molecular dynamics (MD) simulations of the peptide in lipid membranes

Classical MD simulations were performed for five systems characterized in the Supplementary Table 3. All systems were prepared based on previously equilibrated DOPC and cholesterol bilayer consisting of 128 DOPC (64 in each leaflet) and 40 cholesterol molecules hydrated with approximately ~6700 water molecules. LW21 peptide molecule (KKWWLLLLLLLLALLLLLLLLWWKK) was incorporated both in pure DOPC and DOPC with cholesterol membranes in the transbilayer orientation employing a variant of the method advised by Javanainen and Martinez-Seara.^12^ For comparison, the system (DOPC/CHOL/LW21(long)) containing DOPC, cholesterol, and a variant of LW21 (used for the experimental data) with extra flanking residues (GLLDSKKWWLLLLLLLLALLLLLLLLWWKKFSRS) was also simulated (Supplementary Fig. 10). The LW21 molecule was prepared based on an ideally helical structure employing the pdb2gmx tool from the Gromacs software package.^13^ The Amber03 force field was employed for the peptide.^14^ Lipid molecules were described using the Slipids force field^15^ while the `TIP3P` model was employed for water.^16^ Note that the recently developed `Slipids` force field was designed to be compatible with the Amber parameterization. Each system was simulated employing a rectangular prismatic shape simulation box of the size of approximately 6.8 × 6.8 × 8.5 nm with periodic boundary conditions. The temperature of 293 K was controlled employing the velocity rescale algorithm with the coupling constant of 0.5 ps.^17^ The pressure was set to 1.013 bar and controlled in a semi-isotropic scheme using the Parrinello-Rahman barostat algorithm with a coupling constant of 10 ps.^17^ The cut-off of 1.4 nm was used for both van der Waals and short-range Coulomb interactions whereas long-range electrostatics was accounted for by employing the Particle Mesh Ewald method.^18^ All systems were simulated for at least 500 ns with the initial 300 ns treated as equilibration period and hence not used for further analysis. System equilibration was controlled by means of the standard convergence criteria for energy and temperature as well as by convergence of the area per lipid and by ensuring that bilayer properties (for instance, contacts of water and lipids with the peptide) are converged and similar in both bilayer leaflets. Equations of motion were integrated with 2 fs time step. All MD simulations were performed using the Gromacs 4.6.1 software suit and molecular visualization was done employing the VMD package.^13,19^ Standard Gromacs tools together with in-house Python scripts were used for the analysis and visualization of the data with `NumPy` and `matplotlib` libraries being involved. ^20,21^ The ImageJ package was employed for the visualization of lateral densities.

<u>Lateral diffusion maps</u> were calculated independently for each bilayer leaflet by centering (by the data postprocessing) the amino acids that reside in proximity of the considered leaflet and then tracing the diffusion of lipids over an assumed time interval. The interval of 100 ns was chosen for the presentation purposes as it corresponds to a relatively long-time diffusion and the obtained statistics is good for the 500 ns-long trajectories calculated here. Qualitatively the same results were obtained for 50 ns-long intervals.

For quantitative assessment of lipid mobility we deliberately did not use mean square displacement and diffusion constants because we found a consistent estimation of these quantities in systems with and without peptide problematic. More specifically, MD simulations typically suffer from artificial drift of the center of mass of the studied system. This problem is even more pronounced for lipid bilayers as centers of mass of both leaflets usually drift laterally. A simple removal of such motion, independently for each leaflet, is a remedy even though not a perfect one. Of note, leaflets of lipid bilayers were experimentally shown to laterally slide with respect to each other and hence complete removal of their translational motion is not fully correct. In systems with a transbilayer peptide which couples both leaflets, it prohibits to some extent the lateral movement of leaflets with respect to each other but not the lateral movement of the bilayer as a whole. Overall, these differences between simulations of bilayers with and without transmembrane peptide make it very difficult to estimate and, in particular, quantitatively compare lateral diffusion under these two different conditions. Hence, we use autocorrelation function of lipid-lipid contact dynamics to access lipid mobility. This method provides values which are comparable between various simulated systems and does not suffer from the translational motion artifacts. Contact autocorrelation function C(t) is defined as:

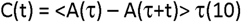

where the function A(t) gives the number of the considered interatomic contacts at time t and the averaging goes over time origins τ in the MD trajectory. The integral of the autocorrelation function over time is the contact correlation time. The decay of C(t) is related to the persistence of interatomic contacts, i.e., slowly-decaying contact autocorrelation function indicates longer-lasting atom-atom contacts. Here, C(t) was calculated employing the “-ac” option of the `g_hbond` Gromacs tool.

<u>The average deuterium order parameters</u>. S_CD_, were calculated in both leaflets as arithmetic average of individual deuterium order parameters estimated for each carbon atom in sn-2 chains of DOPC using the standard `g_order` tool of Gromacs with the *“-d z”* option. Unsaturation of the two carbons in each chain was taken into account. For each carbon atom of a lipid acyl chain, the order parameter is calculated according to the equation:

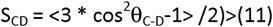

where θ_C-D_ is the angle between the bilayer normal and C-H bonds formed by the considered carbon atom. This parameter is standardly used for the characterization of membrane acyl chains in MD simulations, and it can be directly compared with the deuterium order parameter obtained from ^2^H-NMR data (for more details see ref.^22^). The deuterium order parameter increases together with increasing acyl chain order.

<u>Selection of annular and non-annular lipids</u> was performed using the `g_select` tool of Gromacs. These two groups were defined employing the criterion of 0.75 nm lateral distance between any atom of a lipid and the peptide. The error of the average order parameter was estimated as the arithmetic average of errors (standard deviations) obtained for individual carbons using block averaging.

<u>Radial distribution function</u> (pair correlation function), g(r), gives a probability of finding the considered pair of molecules at the given distance r. Here, g(r) was calculated employing g_rdf Gromacs tool with the option “-rdf mol_com” to take into account distances between centers of masses of the considered molecules.

The error of the estimated time averaged values in MD simulations was calculated employing the block averaging method which accounts for fluctuations and autocorrelations while calculating the equilibrium averages.^23^ The Gromacs tool `g_analyze` with the “-ee” option was employed for this analysis.

### Coarse grain molecular dynamics (CG-MD) simulations of the toy models of cylinder-like structures in lipid membranes

The toy models of cylinder-like structures in lipid membranes were prepared and simulated employing MARTINI force field.^24^ A series of models with varying surface roughness were constructed. To this end, seventy MARTINI beads were packed into a space region corresponding to a cylinder with a diameter of 1.2 nm and a length of 4 nm using PACKMOL code.^25^ This diameter equals to the diameter of helical peptide with an average backbone while the length was selected to allow the domain to span across DOPC lipid bilayer. The number of beads to be packed was chosen in order to obtain ~0.5 nm average distance between individual beads (approximately equal to the minimum inter-bead distance between the considered MARTINI beads). The packing resulted in a cylinder-like object comprising of closely packed MARTINI beads (model M1, see Fig. 1c). Nine beads at both ends of the cylindrical domain were designed as polar, using P4 MARTINI bead type. The remaining 52 beads were designed as nonpolar, using C1 MARTINI bead type, the same as standardly used for nonpolar acyl chains of lipids. The resulting model of transmembrane domain had ~0.5 nm-long polar termini and 3 nm-long nonpolar central section; thus resembling a typical short transmembrane helical peptide, such as LW21. Two toy models with increased surface roughness were generated by removal of randomly chosen 10 (model M2) and 26 (model M3) nonpolar beads (see Fig. 1c). The elastic network method was employed with the elastic bond force constant set to 500 kJ mol^−1^ nm^−2^ and lower and upper elastic bond cut-off to 0.5 and 5.0 nm, respectively to preserve the structure of the toy models during MD simulations.

Simulations were performed with GROMACS 5.0.7 software package with standard parameters advised for MARTINI simulations.^26^ Shortly, 1.1 nm cut-off was used for non-bonded interactions employing the potential-shift-Verlet method. The reaction-field algorithm was applied to account for long-range electrostatics with the relative electrostatic screening parameter of 15. Equations of motions were integrated with 10 fs time step. The temperature of 293 K was maintained using the velocity rescale algorithm with 1.0 ps coupling parameter and the pressure of 1 bar was controlled by the semi-isotropic Parrinello-Rahman thermostat with the coupling constant of 12. Trajectories of 1000 ns were calculated with first 200 ns of each simulation treated as equilibration. The toy models M1-M3 (Fig. 1c) were embedded in a transmembrane orientation in previously equilibrated bilayers consisting of either 256 DOPC molecules (128 in each leaflet) or 256 DOPC with 80 molecules of cholesterol. The method of Javanainen was used for the embedding.^27^ The autocorrelation function of contacts between the beads of lipid tails and the surface of M1-M3 toy models were calculated in the same way as in the case of all-atom MD simulations for membranes without (Fig. 1d) and with (Supplementary Fig. 11).

